# Clamping of DNA shuts the condensin neck gate

**DOI:** 10.1101/2021.10.29.466484

**Authors:** Byung-Gil Lee, James Rhodes, Jan Löwe

## Abstract

Condensin is a Structural Maintenance of Chromosomes (SMC) complex needed for the compaction of DNA into chromatids during mitosis. Lengthwise DNA compaction by condensin is facilitated by ATPase-driven loop extrusion, a process that is believed to be the fundamental activity of most, if not all SMC complexes. In order to obtain molecular insights, we obtained cryo-EM structures of yeast condensin in the presence of a slowly-hydrolysable ATP analogue and linear, as well as circular DNAs. The DNAs were shown to be “clamped” between the engaged heterodimeric SMC ATPase heads and the Ycs4 subunit, in a manner similar to previously reported DNA-bound SMC complex structures. Ycg1, the other non-SMC subunit was only flexibly bound to the complex, while also binding DNA tightly, and often remaining at a distance from the head module. In the clamped state, the DNA is encircled, or topologically entrapped, by the kleisin Brnl and the two engaged head domains of Smc2 and Smc4, and this tripartite ring is closed at all interfaces, including at the neck of Smc2. We show that the neck gate opens upon head engagement in the absence of DNA, but it remains shut when DNA is present. Our work demonstrates that condensin and other SMC complexes go through similar conformations of the head modules during their ATPase cycle. In contrast, the behaviour of the Ycg1 subunit in the condensin complex might indicate differences in the implementation of the extrusion reactions and our findings will constrain further mechanistic models of loop extrusion by SMC complexes.

**SIGNIFICANCE STATEMENT:** DNA needs to be compacted dramatically to fit into nuclei and during cell division, when dense chromatids are formed for their mechanical segregation, a process that depends on the protein complex condensin. It forms and enlarges loops in DNA through loop extrusion. Our work resolves the atomic structure of a DNA-bound state of condensin in which ATP has not been hydrolysed. The DNA is clamped within a compartment that has been reported previously in other SMC complexes, including Rad50, cohesin and MukBEF. With the caveat of important differences that we also uncovered, it means that all SMC complexes cycle through at least some similar states and undergo similar conformational changes in their head modules, while hydrolysing ATP and translocating DNA.

## INTRODUCTION

Structural maintenance of chromosomes (SMC) complexes are essential drivers of chromosome dynamics in all domains of life. In eukaryotes, condensin organises DNA into rod-shaped chromatids during mitosis, cohesin mediates sister chromatid cohesion and interphase chromosomal organisation, and Smc5/6 is involved in DNA repair. In bacteria, SMC-ScpAB and MukBEF promote chromosome segregation by individualising replicated chromosomes (Yatskevich et al., 2019).

In spite of these divergent high-level functions, SMC complexes share a common architecture, making it likely that they also function through common mechanisms. Several SMC complexes have been shown to enlarge DNA loops by a process known as loop extrusion, which is powered by their ATP Binding Cassette (ABC)-type ATPases (Fudenberg et al., 2017). While loop extrusion by yeast condensin and human cohesin has been reconstituted in vitro (Davidson et al., 2019; Ganji et al., 2018; Kim et al., 2019), how they convert the energy from ATP binding and hydrolysis into movement along DNA is not yet clear.

The core SMC complex is a heterotrimeric ring consisting of two SMC proteins and a kleisin, and in condensin these are Smc2, Smc4 and Brn1, respectively (Fig. 1A) (Hirano et al., 1997). SMC proteins are highly elongated when fully extended, with a globular “hinge” domain at one apex and an ABC-type ATPase “head” domain at the other, separated by 50 nm anti-parallel coiled coil domains. The two SMC proteins, Smc2 and Smc4 come together stably via the hinge domains, forming heterodimers. The Brn1 N-terminal domain binds to the head proximal coiled coil of Smc2, the “neck”, and the C-terminal domain to the Smc4 head at a site called the “cap”, thereby creating a closed tripartite ring. To complete condensin, two HEAT repeat-containing proteins Associated With Kleisins (HAWKs), Ycs4 and Ycg1, stably bind to central regions of Brn1.

**Figure 1.**
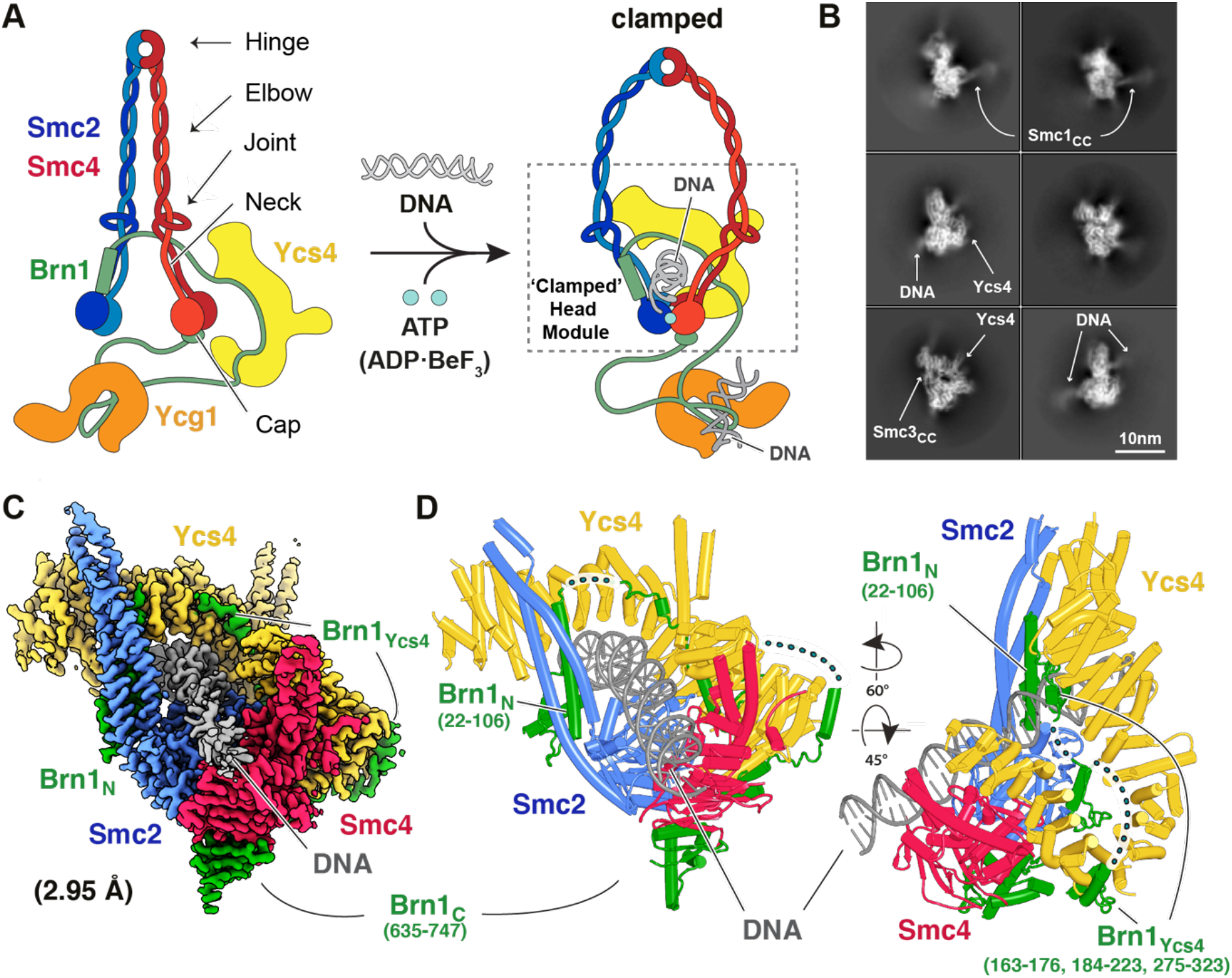
A) Architecture of the yeast condensin SMC complex. Adding DNA and an ATP analogue leads to the formation of the clamped state, which is the subject of this study. The dotted rectangle marks the portion of the head module in the clamped state, of which the cryo-EM structures were solved. B) 2D class averages of cryo-EM images of the condensin tetramer, containing subunits Smc2, Smc4, kleisin Brn1 and Ycs4. The dsDNA is clearly visible, whereas the coiled coil arms of Smc2 and Smc4 and the hinge domains are highly flexible with respect to the well-resolved head module. C) 2.95 Å resolution cryo-EM map of the clamped condensin head module bound to DNA (form I). D) Cartoon representation of the atomic model built into, and refined against the map shown in C. Various parts of the kleisin Brn1 are disordered.

Previous electron cryomicroscopy (cryo-EM) structures of condensin showed that condensin switches between at least two states (Lee et al., 2020). In the absence of ATP, Ycs4 bound to the head domains of Smc2 and Smc4, while in contrast Ycg1 was mobile, presumably connected to the complex by a flexible region within Brn1. In the ATP-engaged structure, Ycs4 unbound and was mobile, whereas Ycg1 bound to the heads instead. Notably, the neck interface between Smc2 and Brn1, where the N-terminal portion of Brn1 binds to the coiled coil neck of Smc2 near the ATPase head, is closed in the absence of ATP but was not resolved in the ATP bound structure. Biochemical experiments demonstrated that opening of the neck interface between Smc2 and Brn1 is driven by ATP binding (Hassler et al., 2019). A similar behaviour was reported for cohesin, whose dissociation from DNA requires opening of the Smc3-kleisin (Scc1) neck interface, which depends on an accessory protein called Wapl and also ATP binding (Chan et al., 2012; Murayama and Uhlmann, 2015).

*In vivo* cysteine crosslinking analyses of cohesin (Collier et al., 2020; Haering et al., 2008), SMC-ScpAB (Wilhelm et al., 2015) and MukBEF (Bürmann et al., 2021) have been used to reveal topological compartments in SMC complexes in which circular DNA can be entrapped (Fig. S1A). These include the SMC-Kleisin (S-K) compartment and, when the head domains are engaged, the Engaged heads-Kleisin (E-K) and the Engaged heads-SMC (E-S) compartments. Sister chromatid cohesion is mediated by the co-entrapment of sister chromatids within cohesin’s S-K compartment. In contrast, it is not currently clear in which compartment the DNAs reside during loop extrusion, or if in any (Cuylen et al., 2011; Davidson et al., 2019).

Cryo-EM structures of cohesin in the ATP-bound engaged (E) state, and bound to one strand of DNA, revealed that DNA is “clamped” on top of the dimerised SMC ATPase domains by Scc2 and the closed neck of Smc3 (Collier et al., 2020; Higashi et al., 2020; Shi et al., 2020). Analysis of the same cohesin-DNA interaction using cysteine crosslinking in vitro revealed that the DNA is not entrapped within the S-K ring, but passes through both the E-S and E-K compartments (Collier et al., 2020). This strongly suggests that in the clamped state of cohesin the kleisin runs over the top of the DNA and that the DNA has not passed through one of the trimer interfaces to reach S-K. Furthermore, a recent structure of MukBEF in the ATP and DNA-bound state conclusively shows that the clamped DNA passes through the E-K and E-S compartments because the entire kleisin MukF is resolved (Bürmann et al., 2021). Interestingly, in this structure, the v-SMC-kleisin neck interface is open, however, the MatP DNA-unloading protein was also bound.

We aimed to investigate how condensin interacts with DNA in the ATP engaged state and how the DNA interaction might regulate the state of the neck interface. To address this, we started by using cryo-EM to solve the structure of condensin in the presence of DNA, ADP and BeF_3_.

## RESULTS

### Cryo-EM structure of a “clamped” condensin complex

To reveal the structure of the condensin-DNA complex we purified a “tetrameric” budding yeast condensin complex lacking the Ycg1 subunit (Fig. S1B, containing subunits Smc2, Scm4, Brn1 and Ycs4). We mixed the condensin tetramer with 80 bp dsDNA, ADP and beryllium fluoride (BeF_3^-^_), applied the sample to cryo-EM grids and collected images (Fig. S2A). ABC-type ATPases bound to ADP-BeF_3_ often represent the enzyme-substrate complex (Oldham and Chen, 2011a). Due to the apparent flexibility of the SMC coiled coil and hinge parts in our data, we were unable to perform analysis of the entire complex (Fig. S2B), but instead focussed on the head module (Fig. 1B, Fig. S3 left). Following the picking of particles with a focus on the SMC head domains, 702,764 good particles showing densities of SMC heads, Ycs4 and DNA were selected with 2D classifications. Through further processing we reconstructed two cryo-EM maps (Form I and II) at resolutions of 2.95 and 3.05 Å, with 286,294 and 251,999 particles, respectively (Fig. 1C and S2C and D). The maps enabled us to build and refine reliable atomic models from both (Table S1). A comparison of the overall conformations resolved showed strong similarity between the two atomic models, with a RMSD (root mean squared deviation) of 1.2 Å (Fig. 1D and S2E).

In both structures, Form I and II, the SMC head domains and proximal coiled coils, Ycs4 and Brn1 form a compact globular structure, the head module. The majority of the SMC coiled coils and hinge are unresolved, most likely due to flexibility with respect to the resolved parts (Fig. 1B and S2B). In both structures, the DNA is held between the upper surface of the dimerised SMC head domains and a groove in Ycs4 (Fig. 1C). The map allowed us to trace a significant fraction of the Brn1 kleisin subunit (279/754, 37% of residues in Brn1), which binds to the neck region of Smc2, to Ycs4 via an extended interaction surface and the cap of the Smc4 head domain (Fig 1D). The neck interface between Smc2 and the N-terminal domain of the kleisin is closed in our structure, possibly indicating that the DNA-bound clamped conformation prevents ATP-dependent opening of this interface, since it had been reported previously that the Smc2/kleisin neck gate opens upon ATP binding in the absence of DNA (Hassler et al., 2019). Form I and II differ in the DNA density where they are not clamped between Ycs4 and the SMC heads. While the DNA of Form I traverses the entire surface of the SMC heads, Form II shows relatively poor DNA density in the region distal to Ycs4 (Fig. S2C, D and E). Because Form I and II therefore differ only slightly, we describe the structure based on Form I, unless otherwise noted (Fig. 1C and D).

### Structure of ATPase head domains and interaction with DNA

Condensin contains two ATPase sites, with each SMC head domain possessing the Walker A and Walker B motifs from one shared or “sandwiched” active site, and the “ABC signature motif” from the other. In the structure, the head domains are dimerised, with two ADP.BeF_3_ sandwiched in the interface (Fig. 2A). Each active site contains a network of interactions between the ADP.BeF_3_, a Mg^2+^ ion and highly conserved residues in Smc2 and Smc4. Engagement of the SMC head domains produces a pseudo-twofold symmetrical DNA-binding surface on their upper side, which is bordered by the two coiled coils (Fig. 2B). Approximately 18 bp of the DNA traverse this surface, interacting with a large number of mostly positively charged residues (Fig. S4A). Interestingly, the DNA is not exactly centred on the pseudo-twofold axis of the ATPase heads (Fig. 2B), and therefore additionally interacts with the coiled coil Smc2 neck and not with the coiled coil of Smc4 (Fig. 2B). The N-terminal domain of Brn1, bound to the neck of Smc2, is also close to the DNA in our structure.

**Figure 2.**
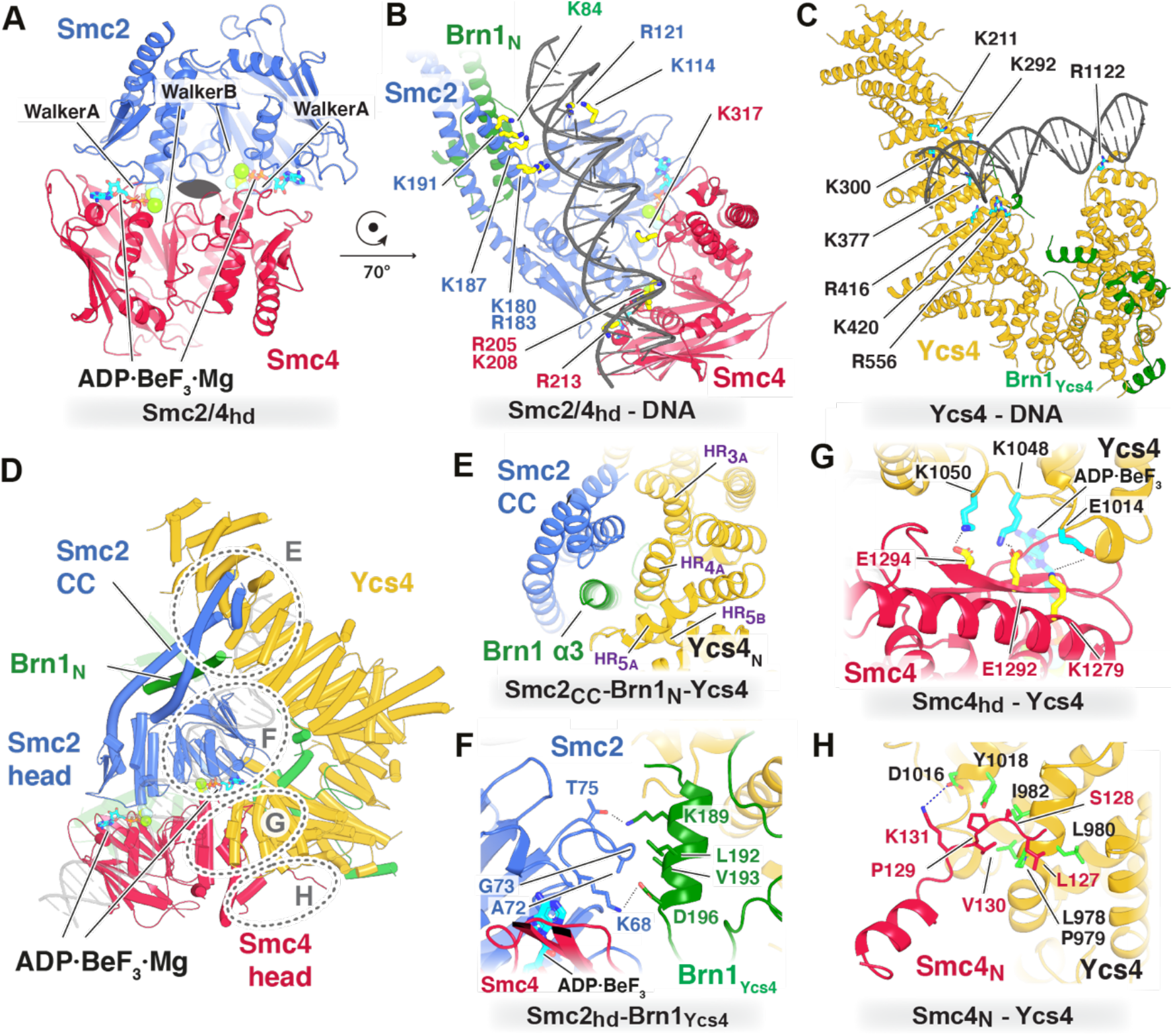
A) Top-view of the pseudo two-fold symmetric ATPase head domains of Smc2 and Smc4. Two molecules of ADP.BeF_3_ are sandwiched between them, leading to the engagement of the SMC heads domains. B) The clamped DNA binds to the largely positively charged upper surface of the ATPase head domains of Smc2/4, but is not perfectly aligned with the pseudo-twofold axis. The DNA also makes contacts with the coiled coil neck of Smc2 and Brn1, but not the neck of Smc4. C) In the clamped head module, Ycs4 makes extensive contacts with the DNA via a positively charged groove towards the N-terminus and through a small contact near the C-terminus. D) Ycs4 make four contacts with other subunits within the head module: E) With the Smc2 neck and Brn1, F) with a loop emanating from the Smc2 head domain that also contacts Brn1 as it binds to Ycs4, G) with a large patch on the back surface of Smc4 and H) with an unstructured and poorly conserved part near the N-terminus of Smc4.

Ycs4 is a large hook-shaped protein consisting of 21 HEAT repeats and a protrusion called the “proboscis” (Hassler et al., 2019) (Fig. S4C). In our cryo-EM map, we observed two clear protein densities running along the inside and around the tip of the hook (Fig. S4C). The first density clearly corresponds to a part of Brn1 that has been identified earlier in the cryo-EM structure of apo condensin and also a Ycs4-Brn1 crystal structure (residues 184-223) (Hassler et al., 2019), while we assigned the second density to a region of Brn1 that had been identified biochemically, but had not been resolved earlier (residues 275-323).

The DNA is clamped onto the top of the dimerised heads by Ycs4. On Ycs4, the DNA binds to a positively-charged surface formed by the edge of a series of HEAT repeats near the middle of the protein, and the surface includes residues K211, K292, K300, K377, R416, K420, R556 (Fig. 2C and S4B). At the other end of Ycs4, and at the other end of the DNA, the tip of the Ycs4 hook also binds the backbone of the DNA through an arginine residue (R1122).

Ycs4 forms extensive contacts with Smc2 and Smc4 at four interfaces (Fig. 2D). The first is a tripartite interaction between Ycs4’s N-terminal HEAT repeats 3 and 4, the neck of Smc2 and the upper end of Brn1’s helix α3 (Fig. 2E). This arrangement traps the Brn1 α3 between Ycs4 and Smc2, possibly preventing opening of the Smc2-Brn1 interface, which, as has been mentioned, has been reported to occur during head engagement in the absence of DNA (Hassler et al., 2019). The second interaction site of Yccs4 is with a loop emanating from the side of the Smc2 head domain (residues 69-73) that also contacts an alpha helix from Brn1 (residues 189-196)) (Fig. 2F). The third interaction is large and is between a region in Ycs4 towards the C-terminus (residues ~1010-1094) and a beta sheet on the back of the Smc4 head domain (Fig. 2G). Finally, an unstructured and not well-conserved region near the N-terminus of Smc4 (127-131) forms the fourth Ycs4 interface, using the outside of the Ycs4 hook (Fig. 2H and S4F).

Having structures of condensin in multiple states (apo, apo-bridged, ATP-clamped) (Lee et al., 2020) (PDB 6YVU, 6YVV and this study) allowed us to analyse the conformational changes that accompany them. The structure of Ycs4 is similar in the apo and clamped structures, but with a 10 Å outward movement of the C-terminal HEAT repeats, which results in a slight widening of the hook (Fig. S4E left). In the apo and apo-bridged conformations, the association between Smc4 and Ycs4 is similar, but the distance between the SMC heads is enlarged in the apo-bridged state, and Smc2 binds to the N terminal HEAT repeats of Ycs4 (Fig. S4E middle). The relative orientation of Ycs4 and the SMC heads is similar. However, in the clamped state there is a 55° rotation of Ycs4 relative to the SMC heads when compared to both apo and apo-bridged (Fig. S4E right).

### In the clamped state, Ycgl binds DNA, but is only loosely attached to the head module of condensin

A previous cryo-EM structure of condensin “pentameric” holocomplex (subunits Smc2, Smc4, Brn1, Ycs4 and Ycg1), showed that in the absence of DNA, Ycg1 binds to Smc2 in the ATP-engaged state (Lee et al., 2020). To determine whether Ycg1 also interacts with the head module in the DNA-bound, clamped state we mixed the condensin pentamer (Fig. S1C) with 80 bp DNA, ADP and BeF_3_ as before, and examined by cryo-EM (Fig. S5A). Using our tetramer structure to guide the analysis, we were able to solve two separate structures after extensive classifications (Fig. S3 right and S5B).

First, we obtained a map of the larger sub-complex comprising the condensin head module, which was almost indistinguishable from the one determined for the tetramer. This 3.7 Å resolution map (processed as a single map due to particle number limitations: 45,112 particles, Fig. S3 right), demonstrated clearly that Ycg1 does not associate rigidly with Smc2, or any other subunit within the globular head module of the complex (Fig. S5B and C).

Second, from the same dataset, we solved the structure of Ycg1 bound to DNA (Fig. 3A and B) at 3.2 Å resolution. In this structure, the region of Brn1 that binds Ycg1 and makes up the “safety belt” (residues 387-524) (Kschonsak et al., 2017) was mostly resolved, with the exception of a disordered connecting loop (residues 411-458) (Fig. 3C). The DNA is somewhat bent as it passes over the DNA-binding surface of Ycg1 and through the safety belt. The DNA covers most of Ycg1’s positively charged surface, and we could identify more Ycg1-DNA interactions than the previous study using X-ray crystallography, which used a shorter DNA (Fig. S5D and E). The fact that we observed Ycg1 bound to Brn1, while not being bound to the head module of condensin strongly suggests that in condensin’s clamped state, Ycg1 remains attached to the rest of the complex via the flexible linkers provided by Brn1 either side of its interaction site with Ycg1 (Fig. 3A).

**Figure 3.**
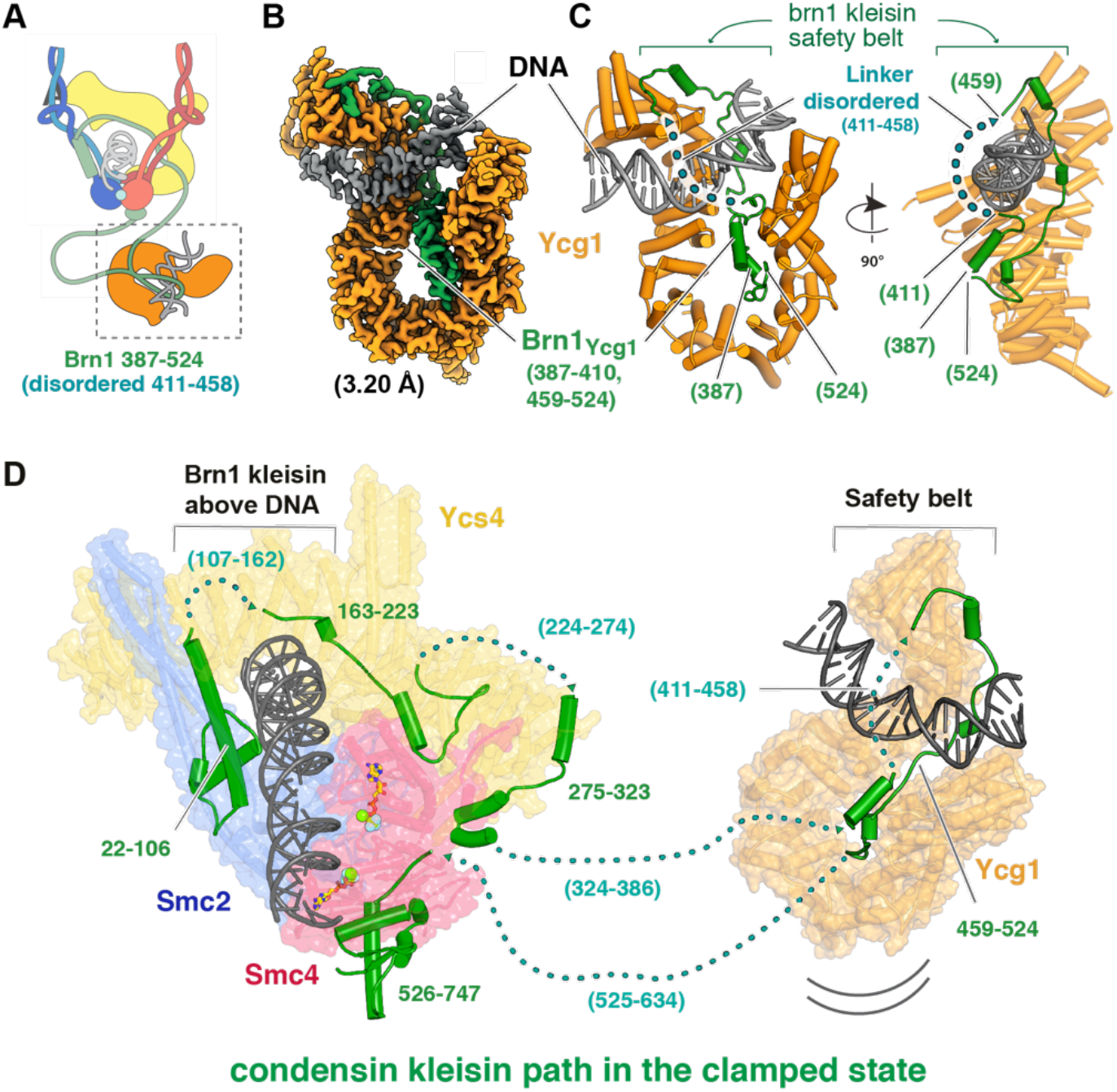
A) Architecture of the condensin pentamer in the clamped state, containing subunits Smc2, Smc4, Brn1, Ycs4 and Ycg1. B) Cryo-EM map at 3.2 Å resolution of Ycg1 alone, bound to DNA, obtained from a dataset collected on the pentamer, presumably because Ycg1 is not rigidly attached to the complex or head module. C) Cartoon representation of the atomic model built into, and refined against the map shown in B. The “safety belt” (Kschonsak et al., 2017) formed by Brn1 is well resolved, except for one disordered loop. D) Proposed path of the kleisin Brn1 through the clamped state of yeast condensin. Several regions could not be resolved in our cryo-EM map (Fig. 1C), but the location of ordered sections suggests the kleisin to remain above the clamped DNA, leading to E-K entrapment. Large Brn1 loops connect Ycg1 to the head module, which enables Ycg1 to reside at quite a distance as shown in Fig. 4A.

### The path of the Brn1 subunit relative to DNA

The path of DNA through SMC complexes is important because it determines whether circular (or very long) DNA is topologically entrapped. In previous clamped structures of cohesin, little of the kleisin subunit Scc1 was resolved and the path was inferred and validated using cysteine crosslinking experiments (Collier et al., 2020). In the present structure, substantially more of the kleisin chain is visible, which helps to determine whether or not the DNA runs through, and is entrapped by, the tripartite ring of Smc2-Smc4-Brn1 (S-K) (Fig. 3D). As described above, in our structure the N-terminal domain of Brn1 (22-106) is bound to the neck of Smc2. Brn1 then snakes along the concave surface of Ycs4 (Brn1: 163-223) and along the C-terminus of the protein (Brn1: 275-323). The next resolved Brn1 residues bind to Ycg1 and form the safety belt around DNA (Brn1: 459-52), before Brn1’s C-terminal winged-helix domain (WHD) binds to the cap of Smc4 (Brn1: 526-747). While several regions remain unresolved, the position of Brn1 residues 106 and 163 indicate that the Brn1 chain most likely passes above the DNA. This means that the DNA runs at least through the E-K compartment (Fig. S1A).

### Interaction between condensin and circular DNA in the clamped state

The condensin-DNA complexes investigated so far in this study were assembled using 80 bp double-stranded linear DNA. Therefore, the clamped head module and Ycg1 were possibly bound to separate DNA molecules and any topological constraints were overridden by the fact that linear DNA was used. To examine the arrangement of the condensin holocomplex when all subunits have the opportunity to bind to the same strand of DNA, we mixed the condensin pentamer, nicked (relaxed) plasmid DNA (1.7 kb), ADP and BeF_3_ and examined the sample using cryo-EM with a Volta phase plate (VPP) to increase imaging contrast (Fig. 4A). In the individual cryo-EM images, without averaging, we observed complexes of condensin where the main body and Ycg1 were most likely bound to the same plasmid. Even without averaging, the plasmid DNA could be seen passing though the head module and also through Ycg1 in the same manner as revealed by our higher resolution structures (Fig. 1C and 3B, respectively).

**Figure 4.**
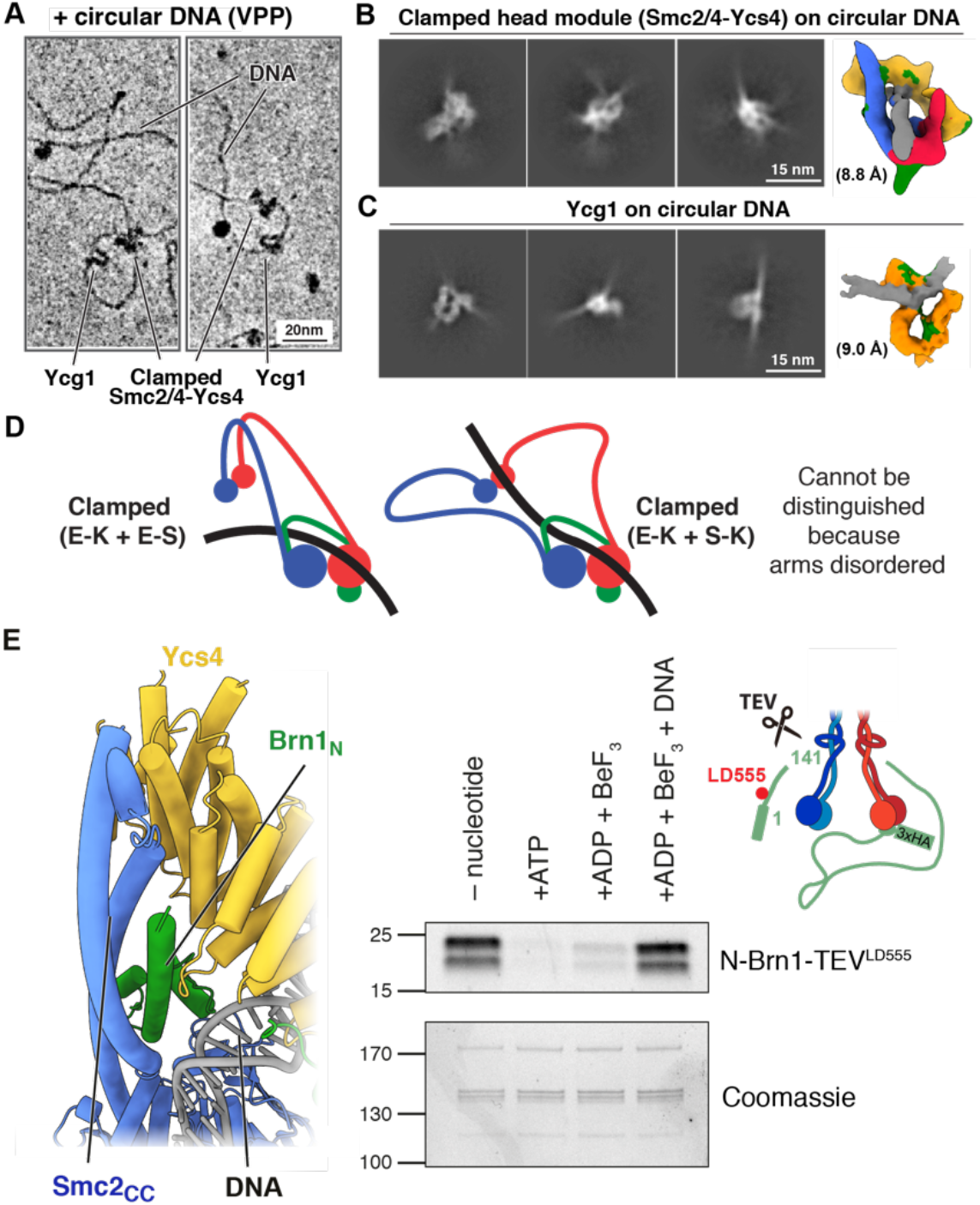
A) Condensin pentamer clamping circular plasmid DNA, as observed by cryo-EM with Volta phase plate (VPP). Ycg1 can be seen at some distance from the head module, presumably because Ycg1 is flexibly attached via Brn1. The SMC coiled coil arms are not resolved. B) The condensin head module clamping circular DNA adopts the same conformation as on linear DNA (compare with Fig. 1C). C) Equally, Ycg1 binds to circular DNA in a manner similar to linear DNA (compare with Fig. 3B). D) The clamped structure of the head module is compatible with two different paths of the DNA with respect to the SMC arms and hinge domains, which are not resolved, leading to different loading paths for topological entrapment. E-K + E-S requires the DNA to pass through the non-engaged heads, whereas E-K + S-K requires opening of a gate in the tripartite S-K ring Smc2-Smc4-Brn1 (see also Fig. S1A). E) Left: the Smc2-Brn1 neck gate is closed in the clamped state of condensin. The neck gate involves the N-terminal helical domain of Brn1 binding to the coiled coil neck of Smc2. An N-terminal portion of Ycs4 and also the clamped DNA may be there to stabilise the neck gate that closes the tripartite S-K ring consisting of Smc2, Smc4 and Brn1. Right: Brn1 N-terminal release assay. Engineered TEV cleavage sites enable the N-terminal Brn1 domain to be cleaved and washed away from beads as long as the neck gate is not closed. In the absence of DNA, addition of ATP or ADP.BeF_3_ open the gate, while the simultaneous presence of DNA keeps the gate shut, presumably through the formation of the clamped state of condensing as shown here.

We then generated 2D class averages and 3D models of the clamped condensin head module and Ycg1, each bound to circular DNA (Fig. 4B and C). While these structures are at lower resolutions due to smaller particle numbers, they showed the same conformations and that the DNA is also entrapped within the E-K chamber. Given that the DNA is circular, it is not possible that the DNA slid into the E-K chamber and therefore there are two possibilities for how the DNA enters. i) One of the three interfaces in the SMC-kleisin ring opens and the DNA is topologically entrapped within this tripartite ring (E-K and S-K) ii) The DNA passes between the disengaged heads before they engage ATP (E-K and E-S). In this second model, the DNA becomes entrapped within both the E-S and E-K chambers simultaneously but never enters the S-K ring. It is important to point out that since the position of the coiled coils and hinge domains could not be resolved in the clamped state, E-S and S-K entrapment cannot be deduced from our head module structure alone (Fig. 4D) and additional topological interactions with either E-S (as deduced for cohesin) (Collier et al., 2020), or with S-K are conceivable. S-K entrapment could be indicated by the opening of the coiled coils in our clamped structure, but it requires gate opening as has been proposed before (Higashi et al., 2020).

### The Smc2-Brnl neck interface remains shut in the clamped state

It has been reported previously that ATP binding, in the absence of DNA results in opening of the Brn1 (N-terminal domain) and Smc2 neck interface (Hassler et al., 2019), and also the equivalent interface between N-Scc1 and the Smc3 neck in cohesin (Chan et al., 2012; Murayama and Uhlmann, 2015). In support of this, in nucleotide-bound and SMC head-engaged structures of cohesin (Muir et al., 2020) and condensin (Lee et al., 2020), both in the absence of DNA, the N-terminal domain of the kleisin is not present, whereas in nucleotide-free and non-engaged structures the kleisin’s NTD is bound (Gligoris et al., 2014; Lee et al., 2020).

Furthermore, a biochemical experiment with recombinant condensin, known as the N-terminal kleisin release assay, has been used to demonstrate that ATP binding opens this interface (Hassler et al., 2019). Recognition sequences for TEV protease were inserted into an unstructured region of Brn1 (after residue 141). After cleavage of Brn1 with TEV protease, in the absence of ATP, the Brn1 N-terminal fragment co-immunoprecipitated (IP) with the rest of the complex, reporting a closed interface. However, when ATP was included during the wash steps, the N-terminal fragment did not co-IP with the complex, indicating that ATP stimulates the disengagement of the interface. Furthermore, the N-terminal fragment was released in a mutant that can bind but not hydrolyse ATP, but was abolished in a mutant that cannot bind ATP. This showed that ATP binding, but not its subsequent hydrolysis, stimulates opening of the Smc2-Brn1 neck interface. Similar experiments have demonstrated that this mechanism is conserved in cohesin at the Smc3-Scc1 neck interface (Murayama and Uhlmann, 2015).

Surprisingly, in our clamped structure the Smc2-Brn1 interface is closed despite engagement of the ATPase heads and nucleotide binding (Fig. 4E left). Two possibilities could explain this finding. i) Unlike ATP, binding of ADP.BeF_3_ does not stimulate release, or, ii) binding of DNA and being in the clamped conformation inhibits release. To test these hypotheses, we used the same N terminal Brn1 release assay (Fig. 4E right). We found that, as previously described, in the absence of nucleotide the N terminal fragment of Brn1 is efficiently co-immunoprecipitated. When ATP was included during the immunoprecipitation wash steps, the N terminal fragment no longer co-immunoprecipitated, confirming that ATP stimulated release. Likewise, when we washed the beads with buffer containing ADP.BeF_3_ the N-terminal fragment did not co-precipitate effectively. This demonstrates that as with ATP, head engagement stimulated by ADP.BeF_3_ results in the disengagement of the Smc2-Brn1 neck interface. In contrast, when the beads were washed in buffer containing both 80 bp dsDNA and ADP.BeF_3_, the N terminal fragment was efficiently co-precipitated. Together, these experiments showed that condensin can adopt two conformations during nucleotide-driven head engagement: in one the Smc2-Brn1 interface is open, as shown in the cryo-EM structure of condensin bound only to ATP or AMP-PNP (Lee et al., 2020), and in the other, reported here, it is firmly shut (Fig. S6A and B). It seems reasonable to state that the only determinant of which conformation it adopts appears to be the presence or absence of DNA.

## DISCUSSION

A mechanism by which condensin transforms interphase chromosomes into mitotic chromatids has recently been proposed: condensin extrudes chromosomal fibres, while hydrolysing ATP, to create an array of DNA loops spanning the entire chromosome (Ganji et al., 2018). However, the molecular basis for this extraordinary motor activity remains enigmatic, despite a significant number of experiments and models (Davidson and Peters, 2021). In this paper we present the structure of condensin in complex with DNA and ADP.BeF_3_. This has revealed several important behaviours of condensin during the ATPase cycle. First, condensin adopts a “clamped” conformation in which DNA is bound between the engaged SMC head domains, the Ycs4 subunit and the Smc2 neck domain. Second, the DNA passes through the E-K chamber between the kleisin and the head domains. Third, the interface between Smc2 and Brn1 is closed. The latter is surprising because head engagement was thought to open this interface (Hassler et al., 2019). We also present biochemical evidence that this ATP-stimulated interface opening is inhibited by the presence of DNA.

The clamped state of condensin is closely related to the clamped state of cohesin (Fig. S6C and D) (Collier et al., 2020; Higashi et al., 2020; Shi et al., 2020), highlighting potential deep mechanistic similarities between the two proteins. In cohesin, the Scc2 cohesin “loader” subunit takes the position of Ycs4 in clamped condensin and both complexes show very similar arrangements around the neck gates (Smc2-Brn1; Smc3-Scc1), with the kleisins’ N-terminal domains wedged in between the SMC neck, the DNA and the clamping subunit. It is therefore not surprising that neck gate opening seems to depend on the absence of DNA and the presence of ATP in both complexes (Hassler et al., 2019; Muir et al., 2020). Because the clamped state of MukBEF is analogous to the clamped state of cohesin (Fig. S6E) (Bürmann et al., 2021), it is also related to condensin and together with the finding that SbcCD (Mre11-Rad50), an SMC-like protein involved in DNA repair also binds DNA in this way, (PDB 6S85) (Käshammer et al., 2019), it seems that DNA clamping is a conserved state of these complexes and is likely involved in their most fundamental functions.

Taken together with published structures (Lee et al., 2020), our data demonstrate that condensin can adopt at least four stable conformations; two very different conformations in the presence of nucleotide (ATP-engaged [Fig. S6A right] and DNA-clamped), depending on the presence of DNA, and the two nucleotide-free apo conformations (apo and apo-bridged [Fig. S4E left and middle]) (Lee et al., 2020). In the apo conformation, Ycs4 is bound to the SMC heads, which are juxtaposed with their ATPase active sites facing away from each other. In the apo-bridged conformation, the heads are forced apart by Ycs4 which sits between them. In both states Ycg1 is bound to the complex only via the flexible Brn1 linkers. In the ATP-bound structure without DNA (ATP-engaged), the heads are engaged but now Ycs4 is no longer bound to the main body whereas Ycg1 tightly binds to the Smc2 head and the Smc2-Brn1 neck interface is open (Fig. S6A right). Finally, in the ATP- and DNA-bound clamped structure, the head domains are engaged around nucleotides, however Ycs4 is associated with the main body rather than Ycg1 - and the Smc2-Brn1 neck interface is closed (this study).

It is conceivable that the different conformations between the ADP.BeF_3_ + DNA and the ATP or AMP-PNP structures (Smc2-Brn1 interface opening and Ycg1/Ycs4 binding to SMC heads) are a result of the different nucleotides that were used to engage the head domains rather than the presence of DNA. This is unlikely for three reasons. Firstly, in previous structural studies of ABC-type ATPases, heads engaged with different nucleotide analogues result in almost identical conformations of these domains (Oldham and Chen, 2011b). Second, in the equivalent clamped structure of cohesin, ATP was used in combination with mutations (EQ Walker B) that result in defective ATP hydrolysis but not binding (Collier et al., 2020). Despite using ATP rather than ADP.BeF_3_, the proteins adopted the same clamped conformation and the Smc3-Scc1 interface was closed. And third, our Brn1 release assay showed no obvious difference between the effects of ATP or ADP.BeF_3_ on neck gate opening.

Our structure raises an interesting question about the significance of the previous ATP bound structure of condensin without DNA. Does condensin adopt two distinct ATP-bound conformations as part of the loop extrusion process? If not, what is the relevance of ATP binding when not in the clamped state? One possible explanation is indicated by the state of the Smc2-Brn1 neck interface in each structure. In the clamped state this interface is shut, whereas in the DNA free structure the Brn1 N-terminal domain cannot be resolved, likely indicating that it is open. This finding is further supported by biochemical evidence showing that ATP binding releases the Brn1 N-terminal domain, unless DNA is included in the reaction. Because opening of this interface compromises the integrity of the tripartite condensin ring, it is possible that the ATP bound state represents a hypothetical unloading reaction. Condensin needs to be able to dissociate from chromatin in a regulated manner and opening of the Smc2-Brn1 interface is one likely mechanism for that release, should loop extrusion or other cellular activities of condensin involve topological entrapment (Cuylen et al., 2011). Indeed, opening of the Smc3-Scc1 interface in cohesin, whose topological entrapment in cells is firmly established (Haering et al., 2008), is similarly driven by ATP binding (Muir et al., 2020; Murayama and Uhlmann, 2015). Finally, a recent structure of MukBEF, in complex with the MatP unloader, which is believed to show the complex poised for topological unloading of DNA shows the analogous N-kleisin neck gate (MukF-MukB) in an open state (Fig. S6B) (Bürmann et al., 2021).

Our cryo-EM data provide clear evidence that condensin can bind to DNA via Ycg1 and the clamped head module simultaneously. Based on our structures and published single molecule experiments (Ganji et al., 2018) we propose that this is an intermediate step in the loop extrusion process and the variable distance of Ycg1 with respect to the head module might determine, or allow condensin to make steps on DNA that are needed to extrude loops and to enlarge them. It seems also worth pointing out that the two binding sites, in the clamped head module and on Ycg1 enable the trapping of pre-formed DNA loops as shown directly in Fig. 4A, and this would be a convenient initial reaction to start loop extrusion.

But it is clear that a precise understanding of the order of the conformations during the loop extrusion cycle will be required to be able to describe the molecular mechanism for DNA translocation and extrusion. This may be achieved using single molecule approaches such as FRET, especially for the presumably very flexible SMC arms (Bauer et al., 2021), but we envisage that resolving these structures of SMC complexes during the loop extrusion process by cryo-EM may ultimately be needed to understand the mechanisms involved at the same level of detail that has been achieved for cytoskeletal or other nucleic acid motor proteins, as long as the important and relevant parts of the complex retain enough rigidity for this to be achievable.

## MATERIALS AND METHODS

### Plasmids and protein expression

Wild type condensin holocomplex (pentamer) was overexpressed in budding yeast from two 2 micron high copy number plasmids (pGAL7 SMC4-3xStrepII pGAL10-SMC2 pGAL1 BRN1-HA3-His_12_ URA3 and pGAL1 YCG1 pGAL10 YCS4 TRP1) (Terakawa et al., 2017). Condensin holocomplex containing TEV cleavable Brn1 was overexpressed from pGAL7 SMC4-3xStrepII pGAL10-SMC2 pGAL1 BRN1(ybbR tag replacing residues 13-23, 3 x TEV site inserted at residue 141)-HA3-His_12_ URA3 and pGAL1 YCG1 pGAL10 YCS4 TRP1.

For expression of the condensin tetramer, YCG1 was deleted by Gibson Assembly to produce pGAL1 YCG1 TRP1. Cultures were grown in -URA -TRP dropout medium + 2% raffinose until OD_600_ of 0.8-1. Protein expression was induced by adding 2% galactose to the medium and incubating overnight at 30ºC.

### Protein purification

Recombinant condensin complexes were purified as previously described, with minor modifications (St-Pierre et al., 2009; Terakawa et al., 2017). Induced yeast cultures were centrifuged, washed once in PBS and centrifuged again. The cell pellet was resuspended in 1x pellet volume of Buffer A (50 mM Tris-HCl pH 8.0, 200 mM NaCl, 5% glycerol, 5 mM 2-mercaptoethanol) with protease inhibitors (Roche) and 300 U/L benzonase (Sigma). The cell suspension was then lysed in a Spex Freezer Mill (5 cycles of 3 min at 12 cpm, with 3 min cooling between cycles). The lysate was clarified by centrifugation for 30 min at 20,000 rpm in JA 25.50 rotor and filtered through Whatman paper. The lysate was then adjusted to pH 8.0 by addition of NaOH. The filtered and clarified lysate was loaded onto two 5 mL His-Trap columns (Cytiva) assembled in series at a flow rate of 1 mL/min. The column was washed with 100 ml of Buffer A + 500 mM NaCl, 50 mL of Buffer A + 40 mM imidazole at 5 mL/min and 50 mL of Buffer A + 60 mM imidazole at 5 mL/min. The protein was then eluted in Buffer A + 200 mM imidazole. Peak fractions were pooled and diluted 2-fold in Buffer SB (50 mM Tris-HCl pH 8, 200 mM NaCl, 5% glycerol, 1 mM DTT) and loaded onto a 5 mL Strep-Trap column (Cytiva) at a flow rate of 1 mL/min. The column was washed with 50 mL Buffer SB, 20 mL Buffer SB with 50 mM KCl, 10 mM MgCl2 and 1 mM ATP (Sigma) followed by 50 mL Buffer SB. The protein was then eluted in buffer SB containing 5 mM desthobiotin (IBA). The peak fractions were pooled and concentrated with a Vivaspin 20 100,000 Da centrifugal concentrator (Sartorius).

For fluorescent labelling of TEV cleavable condensin, at this stage CoA-LD555 (Lumidyne) was conjugated to Brn1-ybbR with recombinant SFP synthase (Addgene Plasmid #75015) as previously described (Yin et al., 2006). All proteins were further purified by size exclusion chromatography on a Superose 6 Increase 10/300 GL column (Cytiva), pre-equilibrated with Buffer SB containing 1 mM MgCl_2_. The peak fractions were concentrated using a Vivaspin 20 100,000 Da centrifugal concentrator and frozen.

### Assembling clamped condensin and Cryo-EM grid preparation

Purified condensin samples were buffer-exchanged using Zeba Micro Spin 7K MWCO columns (Thermo Fisher Scientific) in TNT buffer (30 mM Tris/HCl, 60 mM NaCl, 1 mM TCEP, and 2 mM MgCl2 pH 7.5). Then, 0.5 ~ 1 mg/ml samples were incubated with 1 ~ 2 μM 80 bp dsDNA (5’-GAATTCGGTGCGCATAATGTATAATAAGATAAATAAGCTTAAGTTCTTCCGATGCATAATAACATAATACGTGACTTTAC-3’, and 5’-GTAAAGTCACGTATTATGTTATTATGCATCGGAAGAACTTAAGCTTATTTATCTTATTATACATTATGCGCACCGAATTC-3’, IDT) or 100 nM relaxed circular DNA (1789 bp, derived from pUC19) (Collier et al., 2020) in the presence of 5 mM ADP, 1 mM BeSO_4_ and 10 mM NaF for 30 min at room temperature. The incubated samples were supplemented with 0.1 % (w/v) β-octyl glucoside (Anatrace) and applied onto freshly glow-discharged 200 square mesh Ultrafoil R2/2 gold grids (Quantifoil). The samples were plunge-frozen with a FEI Vitrobot Mark IV (Thermo Fisher Scientific) at 4 °C and 100 % humidity (blotting force −10 to −15, blotting time 1.5 ~ 2 s) and a liquid-ethane cryostat set to 93 K (Russo et al., 2016).

### Cryo-EM image acquisition

All cryo-EM datasets were acquired at 300 kV on a FEI Titan Krios electron microscope (Thermo Fisher Scientific). EPU was used for automatic data collection, and for non-VPP datasets AFIS (aberration-free image shift) was used to increase throughput. For the condensin tetramer:80 bp DNA dataset, images were acquired using a GIF Quantum energy filter with 20 eV slit width and a 100 μm objective aperture. 8,784 movies were recorded using a Gatan K3 summit direct electron detector in super-resolution mode, with a nominal magnification of 81,000x, corresponding to a pixel size of 1.07 Å/pixel (0.535 Å/pixel in super-resolution), and a nominal defocus range of 1.5 ~ 3.3 μm. Each movie was dose-fractionated into 55 frames, with a total dose of 55 e^-^/Å^2^. For the pentamer:80 bp DNA dataset, 4,302 images were collected using a Falcon 4 detector, with a nominal magnification of 75,000x (calibrated pixel size 1.08 Å) and without any objective aperture. The nominal defocus range was set to 1.5 ~ 3.0 μm. Each image was recorded in EER (electron event representation) format with a total dose of 40 e-/Å^2^. For the pentamer:circular DNA dataset, 3,179 images were acquired with a Volta phase plate (VPP) (defocus range: 0.6 ~ 1.1) using a Falcon 4 detector, with a nominal magnification of 75,000x (pixel size = 1.08 Å). Since the number of condensin particles in this VPP dataset was not sufficient to obtain good 3D classes, an additional 1,016 non-VPP images were acquired with a 100 μm objective aperture (defocus range: 1.5 ~ 3.0 μm), using a grid obtained under the same conditions but with a more concentrated sample. Both datasets were recorded in EER format with a total dose of 32 e^-^/Å^2^.

### Cryo-EM image processing

The cryo-EM data processing workflow is summarised in Fig. S3. Processing was performed with RELION 3.1, CtfFind4, crYOLO, and cryoSPARC v3.2 (Punjani et al., 2017; Rohou and Grigorieff, 2015; Scheres, 2012; Wagner et al., 2019), and RELION was used, unless otherwise specified. Overall resolutions were determined based on Fourier shell correlation (FSC) gold-standard criteria (0.143) (Rosenthal and Henderson, 2003). All images were subjected to beam-induced motion correction as implemented in RELION 3.1. Movie frames were aligned and combined with dose weighting using 7 x 5 patches (for K3 datasets) or 5 x 5 patches (for Falcon 4 datasets). The Falcon 4 movies in EER format were dose fractionated into groups of 40 frames, corresponding to a dose of 1 e^-^/Å2 or 0.8 e^-^/Å2 per fractions for the pentamer:80bp DNA or pentamer:circular DNA datasets, respectively. CTF parameters were estimated with CtfFind4.

For the condensin tetramer:80 bp DNA dataset, ~3 M particles were picked with a Laplacian-of-Gaussian blob as template, and subjected to 2D classification. An initial 3D model of the head complex was generated using particles from selected 2D class images showing different orientations. Then, to obtain more particles accurately using crYOLO, a model was trained with the coordinates from the images that formed the selected 2D classes. The resulting ~2 M particles were extracted using a box size of 300^2^ pixels (pixel size=1.07 Å), followed by 2D classification. 702,764 accepted particles were subjected to further 3D classification, and resulted in two 3D classes showing secondary structure features. Further processing was performed separately for each class, but in the same way. First, 3D auto-refinement was performed and resulted in maps of 3.27 Å (Form I) and 3.24 Å (Form II), followed by CTF-refinement for magnification anisotropy, per-particle defocus, per-micrograph astigmatism and beam tilt. For beam tilt and CTF-refinement, the particles were divided into optics groups according to their hole positions (because of AFIS data collection). Then, Bayesian polishing was performed, followed by another round of 3D refinement to generate final maps with 2.95 and 3.05 Å resolutions, for Forms I and II, respectively.

For the condensin pentamer:80 bp DNA dataset, particles were initially picked using crYOLO with a general model. Approximately 600 k particles were picked and a subset of 50 k particles was subjected to a five-class ab initio initial model reconstruction in cryoSPARC v3.2. Using the resulting 5 models, heterogeneous refinement was performed for all picked particles in cryoSPARC. The classes showing features of "clamped" head modules, or Ycg1-DNA complexes were selected, and the corresponding particles for each class were subjected to crYOLO model training separately, followed by automated particle picking. By crYOLO picking with each model, 476,997 particles for the head module and 640,437 particles for Ycg1-DNA were obtained, respectively. The particles for each model were first cleaned-up by cryoSPARC heterogeneous refinement using five models from the first ab initio reconstruction. Additional 2D and 3D classifications without alignment in RELION resulted in 45,112 particles for the head module and 91,024 for Ycg1-DNA. The particles for each were re-extracted with box sizes of 320^2^ and 300^2^ pixels (pixel size=1.08 Å), respectively. 3D auto-refinement performed for each particle followed by CTF-refinement (magnification anisotropy, per-particle defocus, per-micrograph astigmatism and beam tilt) and Bayesian polishing. After 3D refinement, maps at 3.68 Å and 3.20 Å resolutions were obtained for the head complex and Ycg1-DNA, respectively.

For condensin pentamer:circular DNA, particle picking was separately done for the two datasets, VPP and non-VPP, using crYOLO with a general model, follow by extraction with a box size of 360^2^ pixels with 2x binning (180^2^-pixel size, pixel size=2.16 Å). The particles from the two datasets were then merged, an ab initio reconstruction was obtained, followed by multiple rounds of heterogeneous refinement in cryoSPARC. 36,588 and 27,040 particles corresponding to the head module and Ycg1-DNA, respectively were selected, and subjected to 3D homogeneous refinement in cryoSPARC to generate 8.75 and 8.95 Å maps for the head complex and Ycg1-DNA, respectively. During processing of the dataset, we also discovered 2D classes presumed to be SMC coiled coil arms or the hinge (see Fig. S2B), however, we were not able to obtain reliable 3D maps of these parts, presumably due to their flexibility.

### Model building and refinement

For atomic model building, the maps of the two Forms I & II of the head module from the tetramer dataset, and the Ycg1-DNA complex from the pentamer dataset were further improved using DeepEMhancer (Sanchez-Garcia et al., 2021). Model building was carried out in COOT (Emsley et al., 2010)e and ISOLDE (Croll, 2018). Coordinates were refined using phenix.real_space_refine (Afonine et al., 2018). Model validation was performed with MolProbity (Chen et al., 2010). The atomic model for the head module was first built into the 2.95 Å cryo-EM density of Form I from the tetramer:80 bp DNA dataset. The models of Smc2 head-Brn1_25-108_, Smc4 head-Brn1_643-668,685-747_ and Ycs4-Brn1_166-174,184-195,199-220_ were taken from the apo-condensin cryo-EM structure from *S. cerevisiae* (PDB: 6YVU) (Lee et al., 2020), and used as templates. The DNA model was initially taken from the *S. cerevisiae* cohesin cryo-EM structure (PDB: 6ZZ6) (Collier et al., 2020). Each atomic model was docked into the EM map using UCSF Chimera (Pettersen et al., 2004). Coordinates were then manually adjusted and rebuilt in COOT. The DNA model was further refined using the annealing function in ISOLDE. Clear map densities in the vicinity of HEAT repeats 14 to 18 of Ycs4 were apparent after building using the templates, and were identified as Smc4(126-144) and Brn1(275-323), using secondary structure predictions and cryoID (Ho et al., 2020), and then built de novo in COOT. The N- and C-terminal low-resolution regions of Ycs4(26-92 and 1159-1168) were built using ab initio models of *S. cerevisiae* Ycs4 generated by AlphaFold2 (https://alphafold.ebi.ac.uk/entry/Q06156) as a template (Jumper et al., 2021). For Form II of the head module from the tetramer dataset the refined model of Form I was docked into the 3.05 Å map, and rigid-body refined in PHENIX, followed by manual adjustments in COOT. For the Ycg1-DNA atomic model, the crystal structure of the Ycg1-Brn1 complex (PDB: 50QQ) (Kschonsak et al., 2017) was docked into the Ycg1-DNA map from the pentamer dataset in Chimera, and a DNA model was flexibly fitted using ISOLDE. The model was then manually adjusted and rebuilt in COOT, followed by real-space refinement in PHENIX. Figures and movies were generated with PyMOL 2.5 (Schrödinger), UCSF Chimera and ChimeraX (Pettersen et al., 2021). Electrostatic potentials were calculated and displayed in PyMOL.

### Brn1 N-terminal release assay

The N-terminal release assay was performed as previously described with modifications (Hassler et al., 2019). 25 μg condensin holocomplex containing the Brn1 TEVs (after residue 141) and labelled with LD555 (Lumidyne) (1 μg/μl), 1 x acTEV buffer (Thermo), 20 U acTEV protease (Thermo) was incubated at 4°C for 16 hours to cleave Brn1.

For each nucleotide condition, 20 μl of protein A magnetic dynabeads (Thermo) were washed twice in 500 μl wash buffer (50 mM Tris-HCl pH 7.5, 125 mM NaCl, 50 mM KCl, 5 mM MgCl2, 5% glycerol, 1 mM DTT, 0.2 mM PMSF and 0.01% (v/v) Tween-20). The beads were resuspended in 300 μl wash buffer, 3 μg of ant-HA antibody (Roche) was added and mixed at 4ºC for 1 h. The antibody-bound beads were washed two times and resuspended in 300 μl wash buffer. 5 μg of cleaved Brn1-condensin was added to the beads per condition and mixed for 1 h at 4ºC. The beads were washed three times in wash buffer followed by three washes in 300 μl in wash buffer, wash buffer with 1 mM ATP, wash buffer with ADP.BeF_3_ (0.5 mM ADP, 0.5 mM BeSO_4_ and 10 mM NaF) or wash buffer with 80 bp dsDNA and ADP.BeF_3_ (10μM DNA was added and mixed before addition of 0.5 mM ADP, 0.5 mM BeSO4 and 10 mM NaF). The beads were then resuspended in 50 μl 2xSDS (100 mM TRIS-HCl pH 6.8, 4% (w/v) SDS, 20% glycerol (v/v) 0.2% (w/v) bromophenol blue, 0.2 M DTT) buffer and incubated at 65ºC for 5 min to elute the proteins. The eluate was resolved by SDS-PAGE, the retained Brn1 N-terminal fragment was visualised on a Typhoon Scanner (Amersham) and immunoprecipitation of condensin and equal loading was confirmed by Coomassie staining of the eluate.

## ACKNOWLEDGEMENTS AND FUNDING SOURCES

We are grateful to Damien D’Amours (University of Ottawa, Canada), Luis Aragon (MRC London Institute of Medical Sciences, London, UK) and Christian Haering (Universität Würzburg, Germany) for sharing plasmids, yeast strains and technical advice. We would like to thank Nicolas Jean (MRC Laboratory of Molecular Biology [LMB], Cambridge, UK) for a gift of the relaxed plasmid and Yue Liu (MRC LMB) for help with model building. We would also like to thank Dima Chirgadze and Steven W Hardwick (Cambridge University, UK) for assistance with EM data collection and Jake Grimmett and Toby Darling (MRC LMB) for help with scientific computing.

This work was funded by the Wellcome Trust (218621/Z/19/Z to J.R.; 202754/Z/16/Z to J.L) and the Medical Research Council (U105184326 to J.L).

## DATA AVAILABILITY

The model coordinates and cryo-EM density maps were deposited in the Protein Data Bank (PDB) and Electron Microscopy Data Bank (EMDB) with the following accession codes: clamped head module from tetramer dataset, Form I (PDB ID 7Q2X and EMD-13783) and Form II (PDB ID 7Q2Y and EMD-13784); clamped head module from pentamer dataset (EMD-13785) and Ycg1-DNA (PDB ID 7Q2Z and EMD-13786); clamped head module from circular DNA dataset (EMD-13787) and Ycg1-DNA (EMD-13788). See Table S1 for details.

---

**Supplementary Figure S1.**
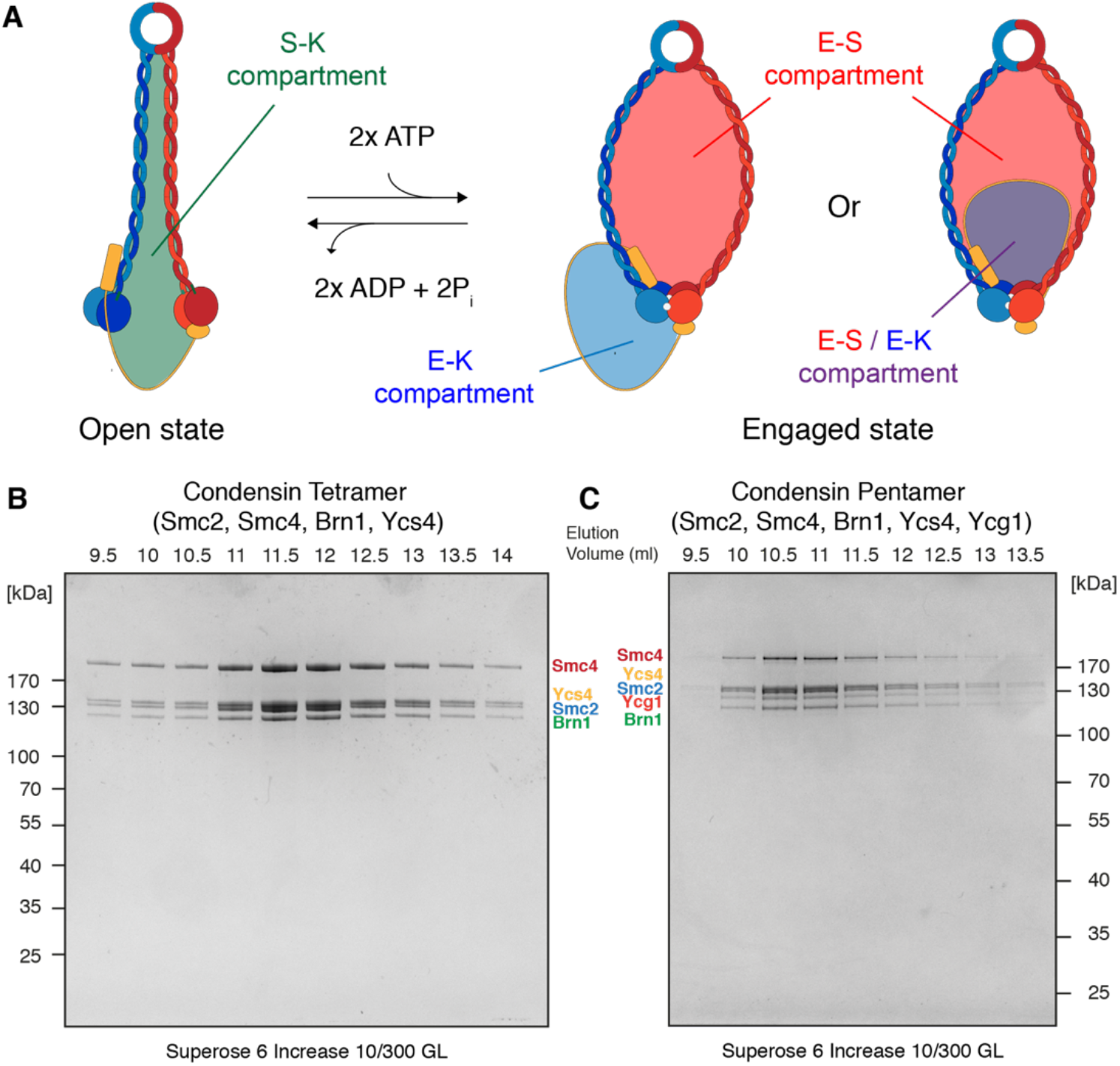
A) Topological compartments of SMC complexes. In condensin, the tripartite Smc2-Smc4-Brn1 ring creates the SMC-Kleisin compartment, S-K. Engagement of the ATPase head domains of Smc2/4 creates two further compartments, Engaged-Kleisin, E-K and Engaged-SMC, E-S. Depending on where the kleisin chain is located, the E-S and E-K compartments can be traversed by the same DNA (right), or not (middle). B) SDS-PAGE gel of the final size-exclusion chromatography of the condensin “tetramer” sample used in this study, comprising subunits Smc2, Smc4, kleisin Brn1 and Ycs4. Elution volumes are provided in mL. C) The same for the condensin “pentamer” sample, comprising Smc2, Smc4, Brn1, Ycs4 and Ycg1.

**Supplementary Figure S2.**
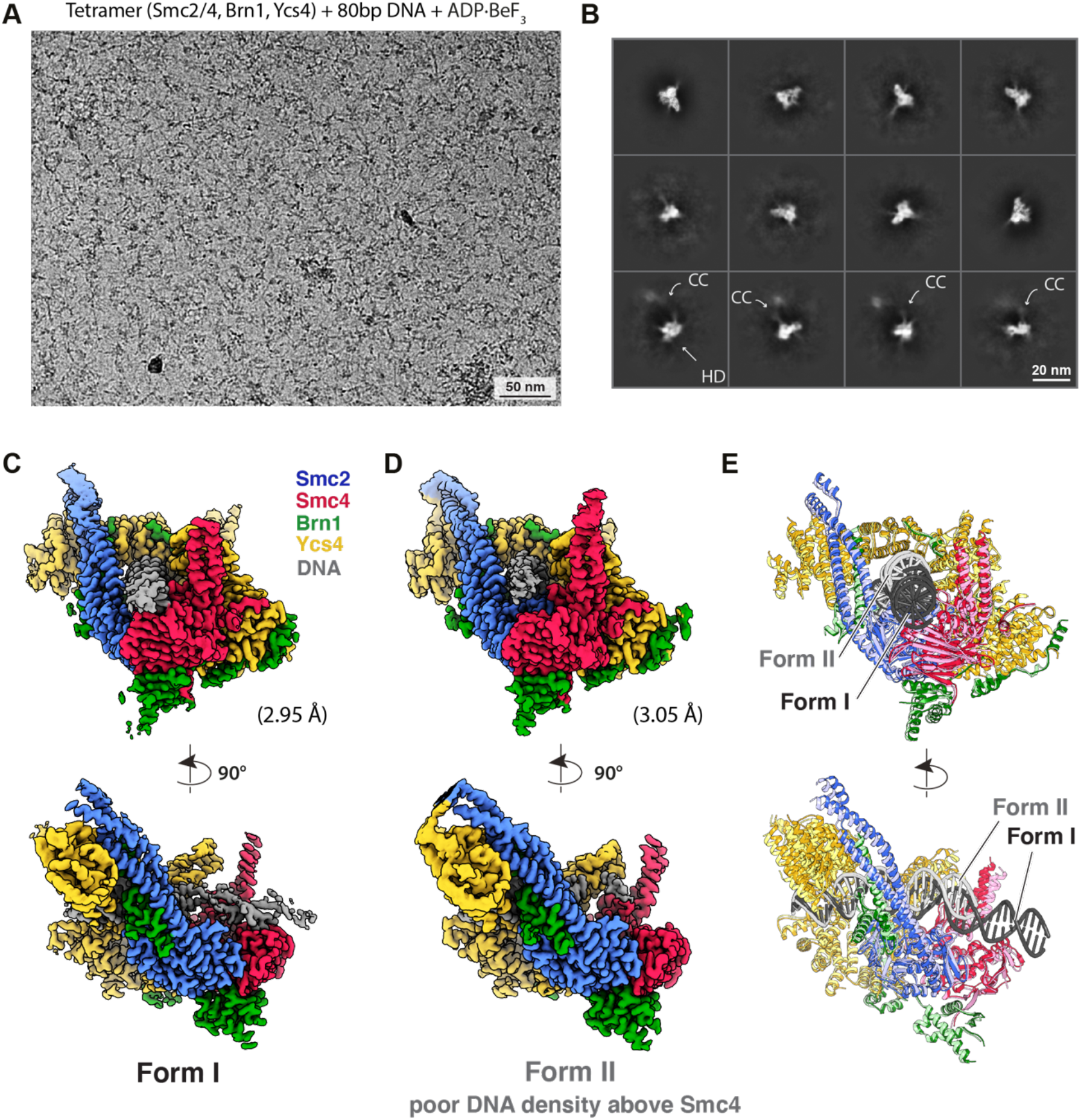
A) Representative micrograph of the cryo-EM dataset used to reconstruct the maps for condensin tetramer in the clamped Form I and II, after adding 80 bp DNA and ADP.BeF_3_, and vitrification. B) A selection of 2D class averages of particles picked from A. The head module is well-resolved and shows secondary structure features (two top rows). The bottom row averages show that most of the coiled coil arms and the hinge domain are flexible with respect to the position of the head module. C) Colour-coded cryo-EM map of the condensin tetramer clamped head module. D) The same for Form II. E) Superposition of the refined atomic models for Form I and II, highlighting the only major difference, the length of the DNA as it extends out of the Smc2-Smc4-Ycs4 clamp above the head domains.

**Supplementary Figure S3.**
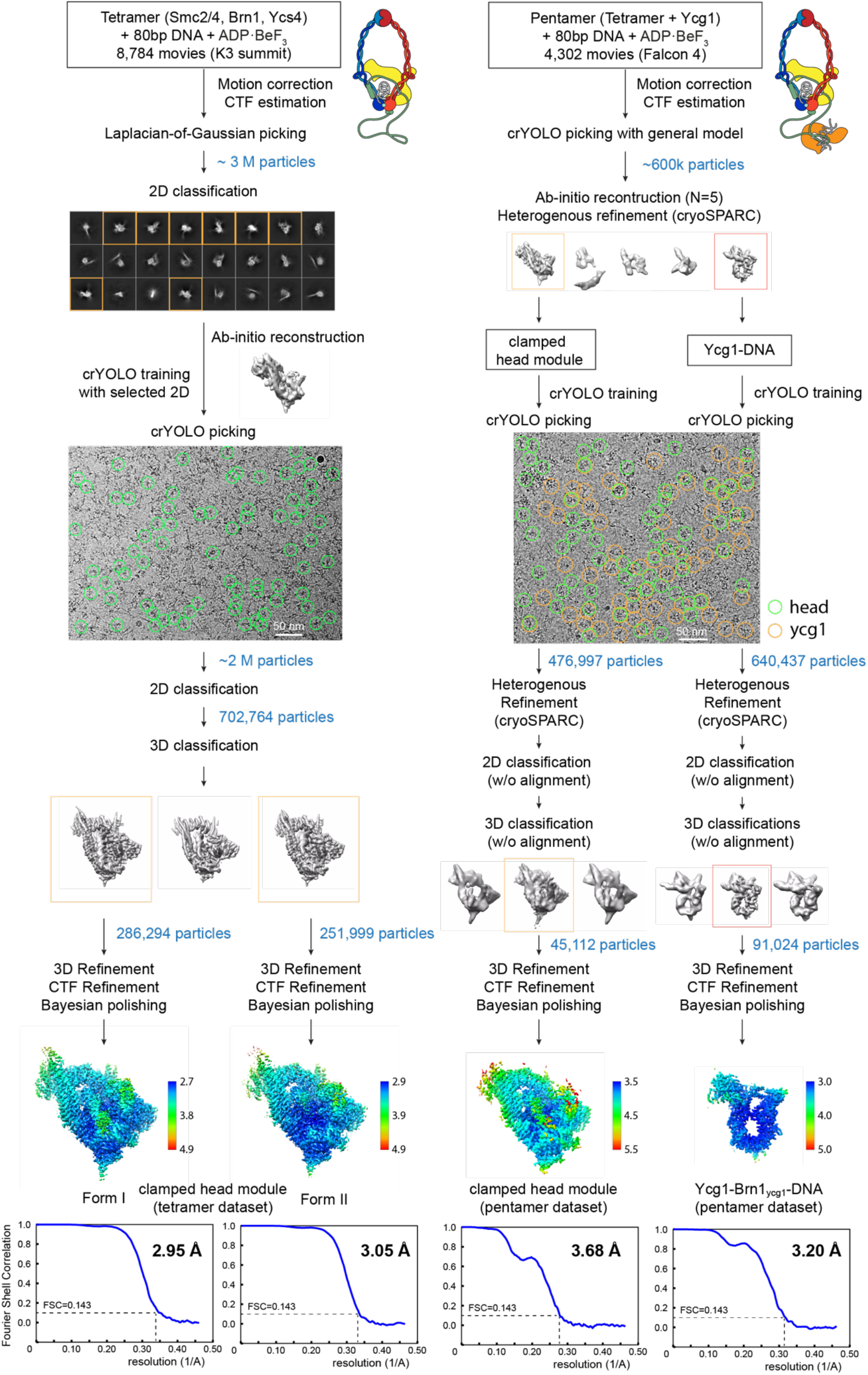
Cryo-EM data analysis and classification workflow. RELION was used unless otherwise specified.

**Supplementary Figure S4.**
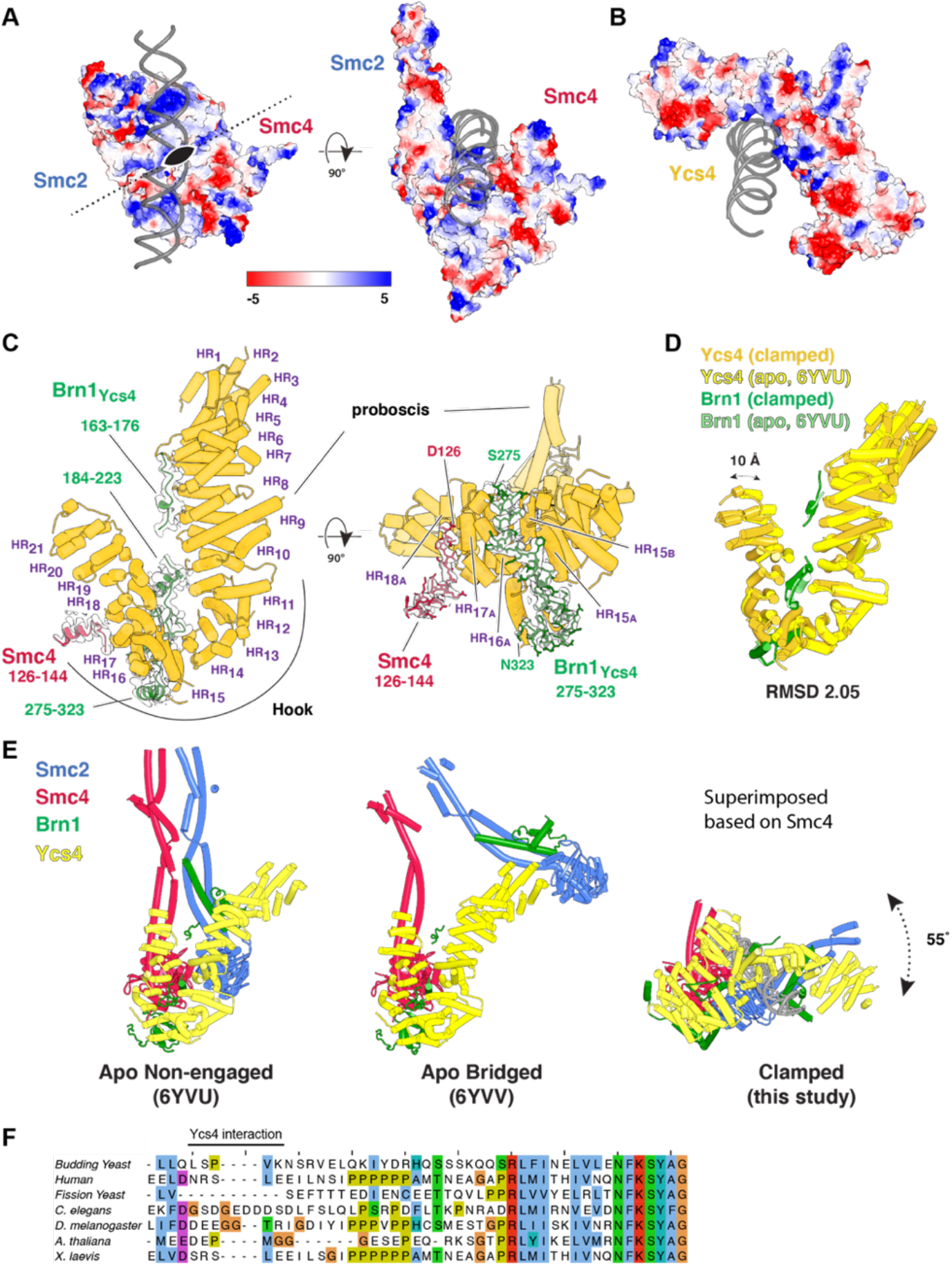
A) Electrostatic potential prediction as calculated in PyMOL, showing positive patches on top of the Smc2/4 head domain heterodimer that coincide with DNA backbone binding. B) The same for Ycs4 as part of the clamp. C) Overview of the HEAT repeat architecture of Ycs4 and showing the cryo-EM map and model of Brn1 sections binding to it in the context of the condensin tetramer clamp. An unstructured part near the N-terminus of Smc4 also interacts with Ycs4 (see also Fig. 2D and H). D) Superposition of the apo condensin Ycs4 structure (PDB 6YVU) (Lee et al., 2020) and Ycs4 as part of the DNA clamp (this study, Form I). A region near the N-terminus moves by up to 10 Å. E) Comparison of condensin tetramer head module structures in the apo (PDB 6YVU), apo-bridged (6YVV) (Lee et al., 2020) and DNA clamped (this study, Form I) conformations. Superposition was done on the head domain of (red) Smc4. Note the models are rotated by 180° around the y-axis relative to Fig. 1C and D (view from the back). F) Multiple sequence alignment of a region of Smc4 (123-169) that includes the Ycs4-interacting region.

**Supplementary Figure S5.**
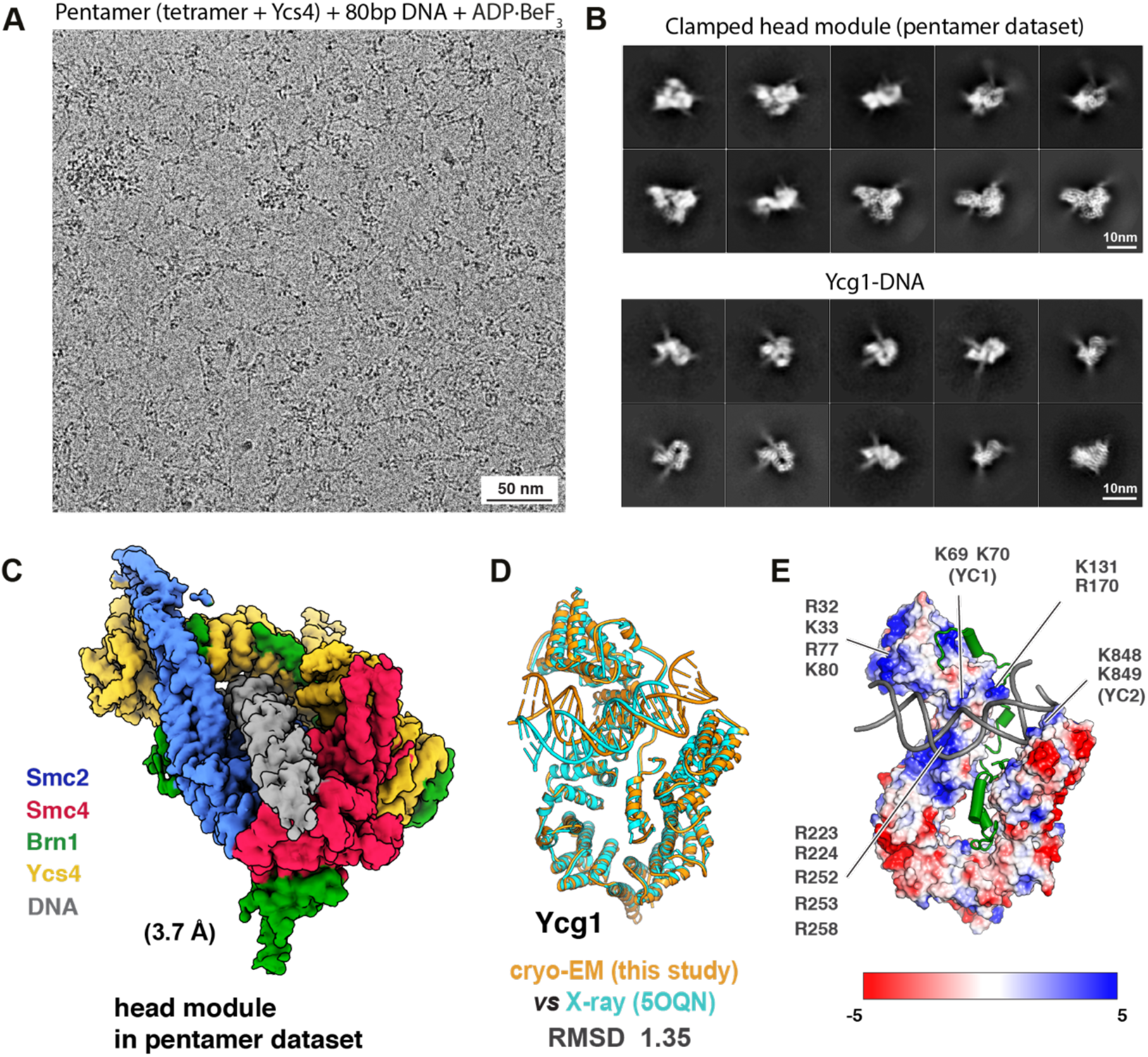
A) Representative micrograph of the cryo-EM dataset used to reconstruct the maps for condensin pentamer in the clamped state, after adding 80 bp DNA and ADP.BeF_3_, and vitrification. B) Top: a selection of 2D class averages of particles picked from A, showing the condensin head module. Bottom: a selection of 2D class averages of particles also picked from A, showing Ycg1 bound to DNA. C) Colour-coded cryo-EM map of the condensin pentamer clamped head module, which closely resembles the tetramer structure (Fig. 1C and S2C and D). D) Superposition of a previous X-ray structure of Ycg1 bound to Brn1 and DNA (PDB 5OQN) (Kschonsak et al., 2017) and Ycg1-DNA as determined here from the condensin pentamer sample. Significant differences in the way the DNA is bound become apparent. E) Electrostatic potential as calculated in PyMOL, plotted onto the surface of Ycg1. Significant positively charged patches are revealed that coincide with DNA backbone binding, including in regions that the X-ray structure did not implicate in DNA binding (the positions YC1 and YC2, which were found in previous work are indicated).

**Supplementary Figure S6.**
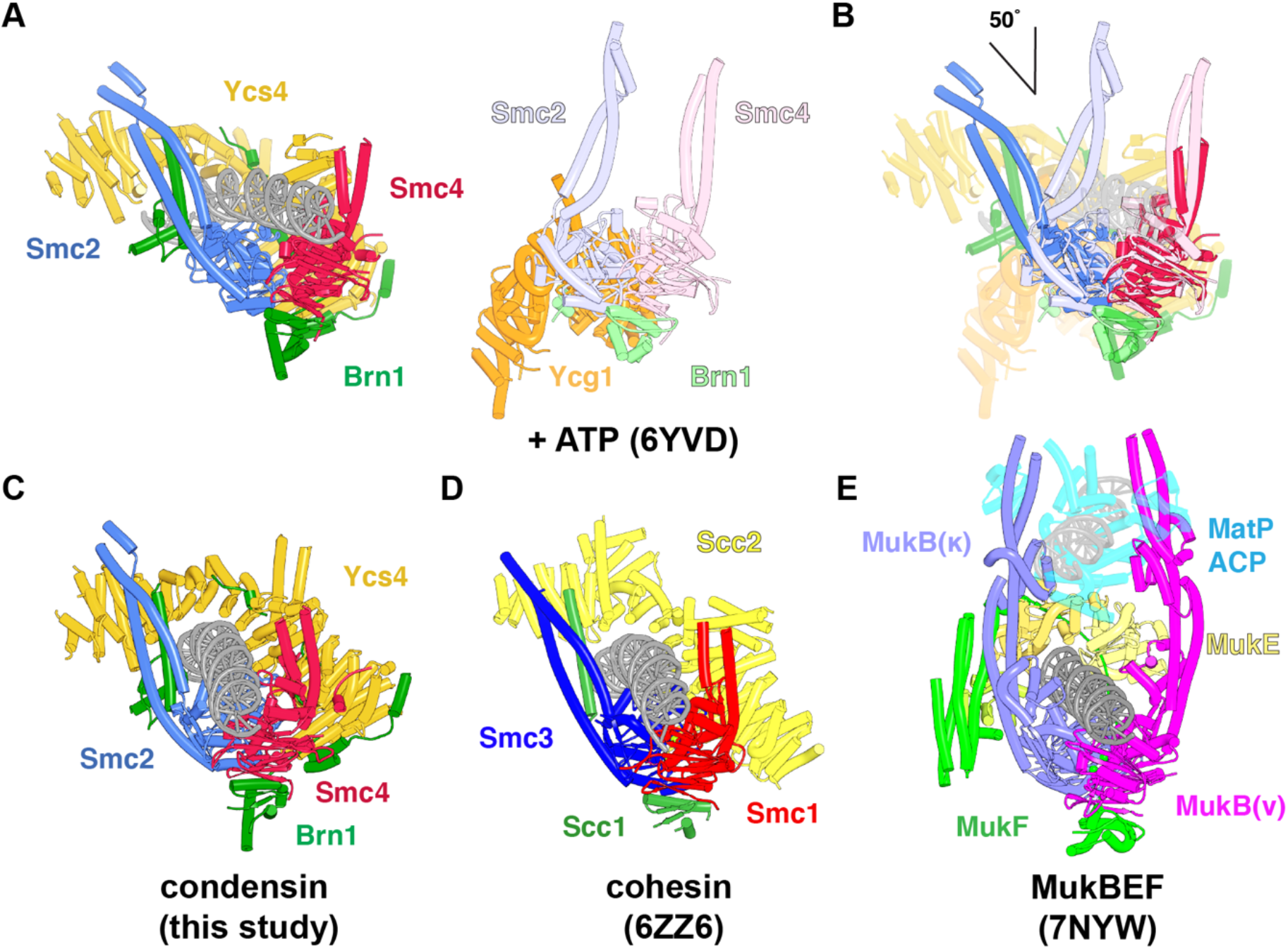
A) Comparison of the DNA-clamped (this study) and DNA-free condensin tetramer structures (PDB 6YVD) (Lee et al., 2020) in the triphosphate, head-engaged states. B) Superposition of A (centred on red/pink Smc4), showing very significant changes in the position of the coiled coil arms of (blue) Smc2 (open in the clamped state, closed by 50° in the apo engaged state). Ycs4 binds to the head module in the clamped state, whereas Ycg1 occupies a similar but rotated position in the ATP-engaged conformation. C, D and E) Comparison of the clamped states of condensin (this study), cohesin (PDB 6ZZ6) (Collier et al., 2020; Higashi et al., 2020; Shi et al., 2020) and MukBEF (PDB 7NYW) (Bürmann et al., 2021). Condensin and cohesin form very similar clamp structures, with Ycs4 and Scc2 creating the same clamp architectures and with the coiled coil arms open. Note that cohesin’s coiled coils remain zipped up in the elbow and hinge regions further up, at least in some of the particles, despite the coiled coil opening near the heads (Collier et al., 2020). MukBEF forms an analogous clamp, with the MukE dimer creating the clamp above the DNA. The kleisin MukF could be traced completely in the structure and was shown to remain above the DNA, meaning the DNA is in the E-K compartment. Note the open neck gate between the helical domain of MukF and the neck of kappa-MukB.

**Table S1.** Cryo-EM data collection, refinement and validation statistics.

|  | Condensin tetramer<br>+ 80 bp DNA |  | Condensin Pentamer<br>+ 80 bp DNA |  | Condensin Pentamer<br>+ circular DNA |  |
| --- | --- | --- | --- | --- | --- | --- |
|  | Head<br>Module<br>(Form I) | Head<br>Module<br>(Form II) | Head<br>Module | Ycg1-DNA | Head<br>Module | Ycg1-DNA |
| **Data collection and processing** |  |  |  |  |  |  |
| Microscope and Camera | Titan Krios, K3 summit |  | Titan Krios, Falcon 4 |  | Titan Krios, Falcon 4 |  |
| Magnification | 81,000x |  | 75,000x |  | 75,000x |  |
| Voltage (kV) | 300 |  | 300 |  | 300 |  |
| No. of micrographs | 8,784 |  | 4,302 |  | 1,016 |  |
| No. of micrographs (VPP) | --- |  | --- |  | 3,179 |  |
| Electron exposure (e<sup>-</sup>/Å<sup>2</sup>) | 55 |  | 40 |  | 32 |  |
| Defocus range (μm) | 1.5 ~ 3.3 |  | 1.5 ~ 3.0 |  | 1.5 ~ 3.0 |  |
|  |  |  |  |  | 0.5 ~ 0.9 (VPP) |  |
| Physical pixel size (Å) | 1.07 |  | 1.08 |  | 1.08 |  |
| Symmetry imposed | *C1* | *C1* | *C1* | *C1* | *C1* | *C1* |
| Initial particle images (no.) | 2,130,610 | 2,130,610 | 476,997 | 694,129 | 105,614 | 105,614 |
| Final particle images (no.) | 286,794 | 251,999 | 45,112 | 91,024 | 36,588 | 27,040 |
| Map resolution (Å) | 2.97 | 3.03 | 3.68 | 3.2 | 8.75 | 9.0 |
| FSC threshold | 0.143 | 0.143 | 0.143 | 0.143 | 0.143 | 0.143 |
| Map resolution range (Å) | 2.6 – 50 | 2.6 – 50 | 3.7 – 50 | 3.0 – 50 | 8.75 – 50 | 9 – 50 |
| **Refinement** |  |  |  |  |  |  |
| Initial model used<br>(PDB code) | Apo-condensin<br>(6YVU) | Apo-condensin<br>(6YVU) | --- | Ycg1 crystal<br>structure<br>(5OQQ) | --- | --- |
| Model resolution (Å) | 3.0 | 3.0 |  | 3.2 |  |  |
| FSC threshold | 0.143 | 0.143 |  | 0.143 |  |  |
| Map sharpening *B* factor<br>(Å<sup>2</sup>) | -52 | -57 |  | -39.4 |  |  |
| **Model composition** |  |  |  |  |  |  |
| Non-hydrogen atoms | 18,240 | 17,755 |  | 8,258 |  |  |
| Protein residues | 2,098 | 2,076 |  | 902 |  |  |
| Nucleotide residues | 72 | 56 |  | 46 |  |  |
| Ligands | 2 ADP<br>2 BeF<sub>3</sub><br>2 Mg | 2 ADP<br>2 BeF<sub>3</sub><br>2 Mg |  | --- |  |  |
| ***B* factors (Å<sup>2</sup>)** |  |  |  |  |  |  |
| Protein | 30.28 | 62.79 |  | 34.99 |  |  |
| Nucleotide | 97.07 | 150.04 |  | 89.42 |  |  |
| Ligand | 22.26 | 54.54 |  | --- |  |  |
| **R.m.s. deviations** |  |  |  |  |  |  |
| Bond lengths (Å) | 0.002 | 0.002 |  | 0.002 |  |  |
| Bond angles (°) | 0.526 | 0.592 |  | 0.482 |  |  |
| **Validation** |  |  |  |  |  |  |
| MolProbity score | 1.53 | 1.71 |  | 1.63 |  |  |
| Clashscore | 7.66 | 8.57 |  | 4.20 |  |  |
| Poor rotamers (%) | 0.00 | 0.00 |  | 0.00 |  |  |
| **Ramachandran plot** |  |  |  |  |  |  |
| Favored (%) | 97.42 | 96.31 |  | 93.36 |  |  |
| Allowed (%) | 2.58 | 3.64 |  | 6.53 |  |  |
| Disallowed (%) | 0.00 | 0.05 |  | 0.11 |  |  |
| PDB ID | **7Q2X** | **7Q2Y** | --- | **7Q2Z** | --- | --- |
| EMDB ID | **13783** | **13784** | **13785** | **13786** | **13787** | **13788** |

